# Homology-based hydrogen bond information improves crystallographic structures in the PDB

**DOI:** 10.1101/147231

**Authors:** Bart van Beusekom, Wouter G. Touw, Mahidhar Tatineni, Sandeep Somani, Gunaretnam Rajagopal, Jinquan Luo, Gary L. Gilliland, Anastassis Perrakis, Robbie P. Joosten

**Affiliations:** Department of Biochemistry, Netherlands Cancer Institute, Plesmanlaan 121, 1066 CX Amsterdam, the Netherlands; San Diego Supercomputer Center, University of California, San Diego, 9500 Gilman Drive, 92093-0505, La Jolla, CA, USA; Discovery Sciences, Janssen R&D LLC, Spring House, PA, USA; Janssen BioTherapeutics, Janssen R&D LLC, Spring House, PA, USA

## Abstract

Crystallographic structure models in the Protein Data Bank (PDB) are optimized against the crystal diffraction data and geometrical restraints. This process of crystallographic refinement typically ignored hydrogen bond (H-bond) distances as a source of information. However, H-bond restraints can improve structures, especially at low resolution where diffraction data are limited. To improve low-resolution structure refinement, we present methods for deriving H-bond information either globally from well-refined high-resolution structures from the PDB-REDO databank, or specifically from on-the-fly constructed sets of homologous high-resolution structures. Refinement incorporating HOmology DErived Restraints (HODER), improves geometrical quality and the fit to the diffraction data for many low-resolution structures. Using approximately 60 years of CPU-time in massively parallel computing, we constructed a new instance of the PDB-REDO databank, a novel resource to help biologists gain insight on protein families or on specific structures, as we demonstrate with examples.

---

Crystallographic structure models are optimized against the crystallographic diffraction data and *a priori* known geometrical observations, the geometrical restraints. We are developing the PDB-REDO procedure, which among many decisions^1^ optimizes the weight between crystallographic and geometrical observations^2^ to re-refine and re-build macromolecular structures before^3^ or after they are submitted to the PDB^4^. In PDB-REDO and any crystallographic refinement procedure however, low resolution diffraction data means that fewer observations of diffracted X-rays are available, and as resolution declines the crystallographic refinement problem becomes increasingly underdetermined^5^. Restraint dictionaries^6,7^ describing ‘ideal’ refinement targets for bond lengths, angles, planar groups and other well-defined chemical features, at low resolution become gradually insufficient to yield high-quality structure models. Additional, external restraints^8^ can be defined and, for example, hydrogen bond restraints^9,10^ (H-bonds) and Ramachandran torsion angle restraints^9,11^ have been used to enhance protein secondary structure quality, particularly at lower resolution.

Macromolecular crystals diffract X-rays to higher or lower resolution in an unpredictable manner: even very similar proteins or the same protein bound to different ligands (e.g. drug candidates), can yield crystallographic data at different resolutions. This allows refinement methods to harvest information from a high-resolution “reference” model and use it to refine low-resolution models^9,10,12–15^. Available implementations of this principle focus on harvesting restraints from a single external reference structure model of high quality, and transferring that information to the low-resolution structure under refinement. Thus the crystallographer is faced with the often difficult and inevitably subjective decision of selecting the ‘best’ model from a group of protein structure models as a reference^16^. Recently, this process was partly automated in the LORESTR pipeline^17^ which uses a series of different refinement protocols and reference restraints from ProSMART^10,12^, to ultimately return the best result using restraints from the optimal reference model.

A set of reference models consisting of many available homologous higher resolution structures, would take conformational flexibility implicitly into account and may therefore help obtaining a better measure for the variation of certain distances, while idiosyncrasies of a single reference model will not cause bad restraint targets. However, heterogeneity in the reference data (e.g. multiple conformational states of a protein) will often be present in the structure ensemble. Therefore, flexibility toward local dissimilarities between the homologs is required. Such flexibility can be achieved by ignoring quasi-random distances or angles and focusing on hydrogen bonds (H-bonds) instead. H-bond networks are well conserved between homologous proteins^18^, and if a certain H-bond is not, inspection of the molecular geometry reveals this immediately. In addition, H-bonds are omnipresent in proteins: more than 90% of all main-chain donors and acceptors are involved in at least one H-bond and side-chain donors and acceptors make more than one H-bond on average^19^. Main-chain H-bonds form the secondary structure elements^18^, and have been restrained in low-resolution refinement before^9,10^. H-bonds that involve side-chains describe the tertiary and quaternary structure of a protein and are therefore more informative about the specific molecular details of a protein.

Here, we present a system that employs H-bond restraints to improve the geometry of low-resolution structure models. First, we optimize targets for H-bond restraints based on global high-resolution structure data from PDB-REDO, and show that these restraints improve protein structure models. Then, we describe how restraint targets can be redefined based on homologous structure data and how both global and homology-based restraints are implemented in the PDB-REDO pipeline. Subsequently we apply our HOmology DErived Restraints (HODER) to the entire PDB data bank, using a highly parallel computational architecture that allowed 60 CPU years of computation to be performed in a few days, allowing a new resource <u>(https://pdb-redo.eu/)</u> to be made publically available. Finally, we present examples of the information that can be derived from this novel resource, and how this can help scientists gain a better understanding of protein structure.

## Results

### Derivation, application and validation of general H-bond restraints

We based the detection of H-bonds on the geometrical criteria defined by McDonald and Thornton^19^, which were slightly loosened to obtain a complete H-bond set (Figure 1A). This set is then subjected to numerous filters to finally arrive at a concise set of high-quality H-bonds that will be restrained. For example, we check each main-chain H-bond against secondary structure information derived from DSSP^20,21^, and donors are not allowed to donate more H-bonds than the number of hydrogen atoms that are bound to them. Not all H-bonds are equal: the distance between the donor and acceptor atom differs between different secondary structure elements and different types of side-chain H-bonds. Therefore, we derived specific targets for each H-bond type from high-quality structural data from PDB-REDO^1^ models with a resolution ≤ 1.8 Å and an R_free_ ≤ 0.20, 10,173 entries in total. H-bonds were detected in all of these entries and separated per category. Main-chain H-bonds were separated in six secondary structure categories (α-helix, π-helix, 3_10_-helix, antiparallel β-strand, parallel β-strand, and others) based on the assignments in DSSP. Side-chain H-bonds were divided into categories where all H-bonds have the same donor and acceptor type. Hence, for example, one category contains all Lys-Nζ to Gln-Oε H-bonds. The full procedure is detailed in the Online Methods.

**Figure 1:**
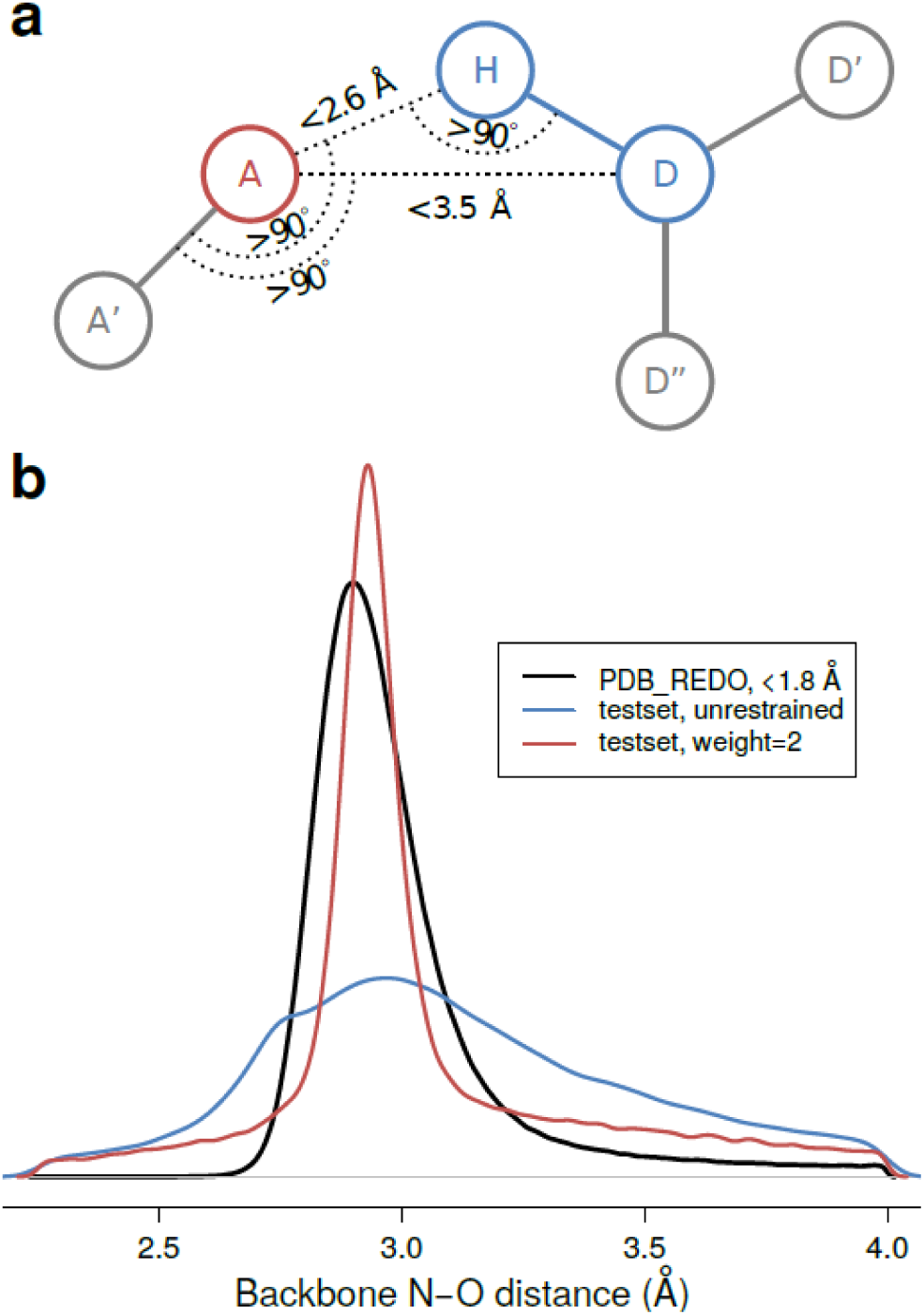
(**a**) Criteria for H-bond geometry. H is the hydrogen atom, D and A the donor and acceptor heavy atoms. Atoms attached to D and A are labelled D’, D” and A’. Each H-bond must fulfill the two distance and three angle criteria indicated. However, if the D-H-A angle is greater than 140°, we allow H-A distances up to 2.75,Å. These criteria were empirically optimized from original work by McDonald and Thornton^19^. (**b**) Distribution of H-bond distances for different restraint weights and groups of structure models. The black line indicates the distribution of all H-bonds in the high-quality PDB-REDO structure model set (resolution < 1.8 Å, and R_free_ < 0.20); the colored lines indicate the distribution in the test set changes after refinement without H-bond restraints (blue) and after refinement with an external restraint weight of 2 (red). The distributions for high-resolution PDB-REDO models and re-refined models from the test set are based on 3.1m and 159k H-bonds, respectively. A restraint weight of 2 makes the test-set distribution resemble the high-resolution set most; other restraint weights are shown in Figure S1.

We detected approximately two million main-chain H-bonds and two million side-chain H-bonds, which were used to derive a target for each H-bond type. The observed H-bond-length distributions were modelled with two-sided normal distribution to obtain ideal target values (see Online Methods). Main-chain targets vary between 2.86 Å and 2.98 Å for different secondary structure elements (Table S1) and side-chain H-bonds between 2.60 Å and 3.36 Å for different types (Table S2). Notably, H-bond restraints previously incorporated into ProSMART/Refmac5^12^ and Phenix^9^, use distance targets of 2.8 Å (Refmac5) and 2.9 Å (Phenix) for all H-bonds, hence the systematic mining from the homogeneously refined structures in PDB-REDO brings new information to the target function.

Defining the weight of H-bond restraints against other restraints during crystallographic refinement is key. This weight was optimized based on the premise that high-resolution structures accurately reflect hydrogen bonding in proteins. Hence, the distribution of H-bond distances was evaluated for the same set of high-quality PDB-REDO models used to derive the targets and also for PDB-REDO entries with a resolution ≥ 2.5 Å. The restraint weight was optimized selecting a value that transformed the H-bond length distribution of the low-resolution set to become most similar to that of the high-resolution set after refinement (Figure 1B, S1).

The effect of the H-bond restraints was initially evaluated by running refinements with and without restraints: the effect of H-bond restraints was greater at lower resolution, while at resolution better than 2.5 Å, the effect of H-bond restraints was negligible. We thus constructed a test set containing 155 low-resolution entries (for details see Online Methods) and proceeded by validating the effect of our method in refinement.

H-bond restraints on the basis of general targets improve the refinement of low-resolution structure models in the majority of cases (Figure S2, Table S3). Mainly the geometry of the protein, measured by packing and Ramachandran angle quality, is improved, while marginal average effects are observed for R_work_ and R_free._ As expected, main-chain H-bond restraints had more impact than side-chain restraints (Table S3). To further test the effect of our new H-bond detection algorithms we repeated calculations with H-bond restraints generated by ProSMART and Phenix: for all model quality criteria our method performs comparably or better than previous methods (Table S4).

Analyzing the general H-bond restraints more specifically showed specific shortcomings: at places the restraints were too tight, distorting the backbone; in some categories specific H-bonds could be relatively weak and should be restrained at greater H-bond length; variation in H-bond length was larger in variable regions such as loops and side-chains; and there are small systematic differences within groups that were assigned a single target (e.g. a systematic difference in H-bond lengths between the middle of a long a-helix and its C-terminus^22^). Because the variability inherent to H-bond lengths cannot be captured in any sensible general division, we set out to define a target on the basis of the homologous structure models, expecting a much more accurate measure of the molecular context of the H-bond than the general data-mining described in this section.

### Homology-based H-bond restraints

To generate homology-based H-bond restraints, we first need to extract the protein sequence from the working PDB file. The program *pdb2fasta* (see Online Methods for details) aims to extract the sequence for modeled and unmodeled parts of the structure, and also maps 73 common types of non-natural (mostly post-translationally modified) amino acids to their parent amino acid. The program has been tried and tested for PDB-wide stability, gives information on unmodeled parts of the sequence, and may therefore be used also for purposes outside the scope of this work.

The sequence file produced by *pdb2fasta* is then passed to BLAST ^23^, which runs it against the PDB-REDO databank^1^ that contains models of a higher average quality^24^ than the PDB^16^. BLAST results are passed to our new program HODER (HOmology DErived Restraints), to first identify suitable homologs from a databank of structural data. Briefly, we consider hits with ≥70% sequence identity and a resolution higher than the query (see Online Methods for details). Importantly, users can also add their own PDB files to HODER, to be used as extra homologs: this functionality is important if one is e.g. working on a series of ligand soaks.

After the residues of the working structure are mapped onto their homologous residues, HODER attempts to derive the H-bond distance restraints. For every H-bond in the working structure, the same H-bond is computed in all homologs, wherever possible. Then, these distances are subjected to 1D *k*-means clustering^25^, the optimal number of clusters is determined by the Bayesian information criterion^26^, within some constraints, and corresponding target distances for each cluster are computed, wherever possible (for details on all the above criteria see Online Methods).

We then repeated the same calculations for H-bond restraints based on general targets for restraints based on homology: as the latter differ from general H-bond restraints only in how their target is derived, the same restraint weight was used. In our test set (see above) 87±16% of the H-bond restraints in each structure were based on homology; for H-bonds where no homology-based target could be defined, we resort to the general target values detailed above. Altogether, homology-based restraints do not deliver a uniform global improvement in performance compared to our general H-bond restraints, but neither did they show obvious drawbacks. Importantly, however, the implementation in PDB-REDO, which we shall discuss now, shows that homology-based restraints work better than general restraints in more extensive model optimization protocols.

### Homology-based H-bond restraints in PDB-REDO

The H-bond restraint procedures have been incorporated into the PDB-REDO pipeline (see Online Methods for details). About one quarter of the crystallographic structures in the PDB (24,506 out of 101,347 PDB-REDO databank entries) with a resolution equal to or worse than 2.5 Å, and thus could benefit from H-bonds restrains. Homology-based restraints could be generated for 17,824 of these entries (73%), and 82% of the total restraints for this set were homology-based (the remaining 18% were defined using the fallback general targets).

In general, the PDB-REDO pipeline already improves both the geometry and the fit to the data of published structure models^1^. When H-bond restraints are used, these improvements are enlarged (Figure 2, Table S5). Importantly, and in contrast to refinements discussed in the previous section, homology-based restraints work decidedly better than general restraints in the PDB-REDO pipeline (see Online Methods for details).

**Figure 2:**
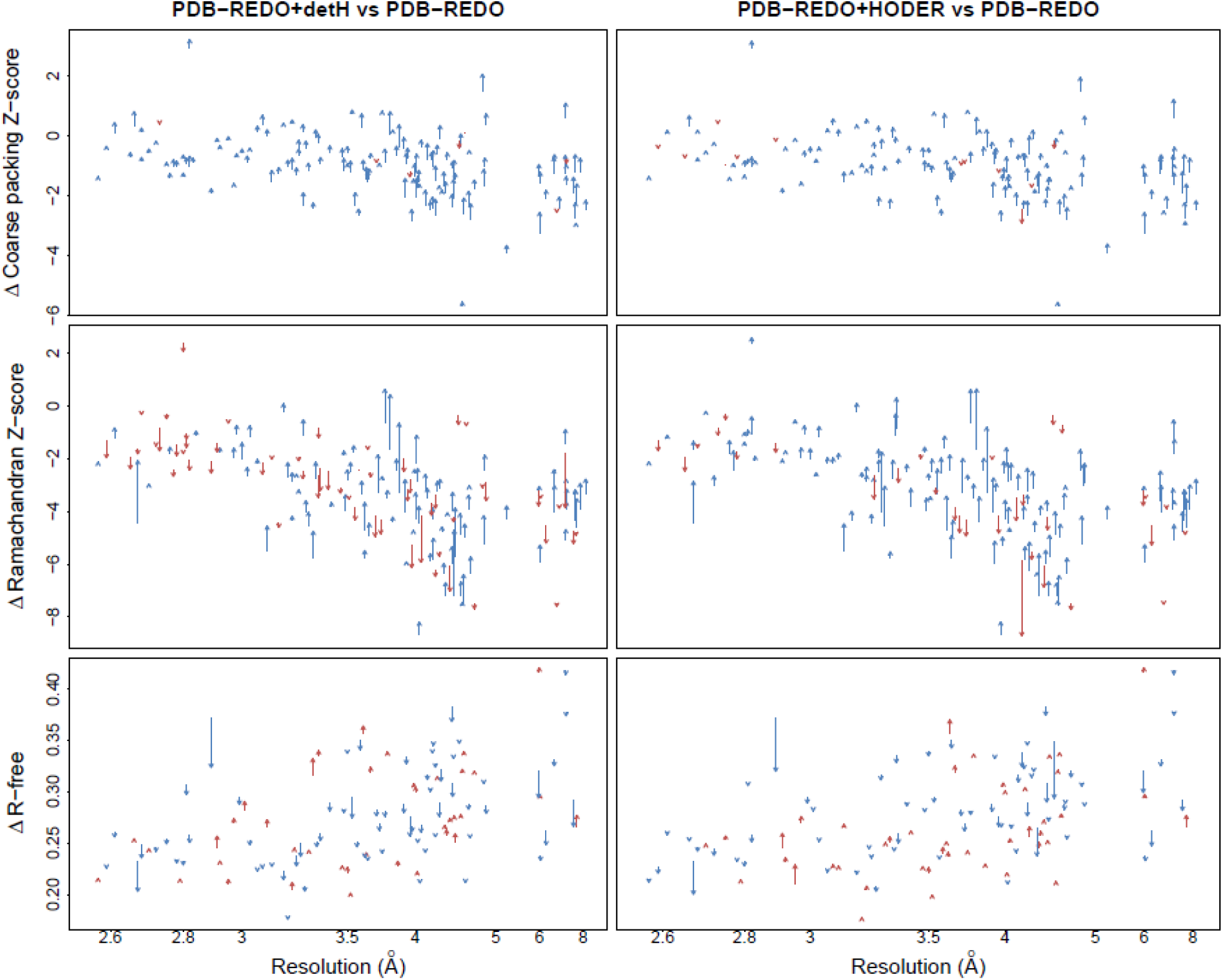
Comparison of PDB-REDO runs with and without general (left) and homology-based H-bond restraints (right) for all entries in the test set. Each arrow represents the scores from two re-refinements on a single PDB entry. Arrow tails indicate scores from refinement without restraints; arrowheads indicate scores from refinement with restraints. Blue and red arrows indicate improvement and deterioration of the score, respectively. The shown scores are the first generation packing Z-score (top) and the Ramachandran Z-score (middle) from WHAT_CHECK^27^ and the R_free_ (bottom) calculated by Refmac5^28^. Arrows at the same resolution have been shifted up to 0.05 Å to reduce clutter. Packing Z-score and Ramachandran Z-score are not shown if they were not computed by WHAT_CHECK; R_free_ is not shown if a new R_free_ set was chosen by PDB-REDO^1^.

When models are subjected to the PDB-REDO pipeline using homology-based restraints, they are influenced by their homologous PDB-REDO entries. In turn, the PDB-REDO models subjected to homology-based restraints may also become the basis of the restraints for other homologous structure models of even lower resolution, which could cause a feedback loop and structure families converging to a consensus structure over multiple rounds of optimization. Then, the true differences between the different structures could be lost. We assessed this risk by subjecting all entries in six protein families (hemoglobin, BRCA1, MutS/MutL, OmpF porin, F1-ATPase and alcohol dehydrogenase) to PDB-REDO five times. Differences between structure models do not decrease when multiple cycles of PDB-REDO with H-bond restraints are applied (Table S6), suggesting that weight optimization and the tolerance to external restraint outliers in Refmac5 prevent bias toward other, possibly incorrect conformations (see Online Methods for details and additional observations).

### Massively parallel computing for a novel PDB-REDO databank with homology information

The observations that global and homology-based H-bond restraints improve low-resolution structure models after a single PDB-REDO refinement encouraged us to update all entries in the PDB-REDO databank^21^ with the most recent version of the (fully automated) PDB-REDO software that includes the refinement strategies based on H-bond restraints.

Based on benchmarking of PDB-REDO jobs, we estimated that we would need a century of single core computing time to calculate all PDB-REDO entries. As this was incompatible with current funding constraints and academic tradition for completion of doctoral thesis work, we decided to deploy PDB-REDO in a High Performance Computing (HPC) environment. Self-contained Docker (www.docker.com) and Singularity^29^ images with all PDB-REDO core and third-party components (more than fifty independent pieces of software) were created to facilitate massive deployment on any (HPC) host (see Online Methods). Running the complete PDB-REDO pipeline with 101.570 entries finally required about 60 CPU years (half a million hours) and all computations were finished within about a week using approximately 3072 cores on the Gordon HPC cluster, and the large memory nodes on the Comet HPC cluster, at the San Diego Supercomputer Centre. These runs were assembled in a new PDB-REDO databank.

This new databank is a resource of consistent and high-quality protein structure models. The introduction of homology-based restraints has improved the quality of low-resolution structure models in a consistent manner, as all low-resolution structure models in the databank were allowed to refine using information derived from high-resolution homologs (Figure 3). Importantly, it is not only a new resource in its own right, but it can also serve for better homology restraint generation for future structure refinement. In addition, the entire PDB-REDO databank is constructed with a single software version (which has not been possible before) and with new algorithms: these include better treatment of twinning, general improvements to TLS, NCS, and ADP refinement^1^, validation and correction of structural zinc sites^30^, better handling of carbohydrates^31^, improved selection of resolution cut-off and the generation of anomalous difference maps when possible. All these developments are consistently and uniformly applied in all entries, in addition to the applicable homology-derived restraints.

**Figure 3:**
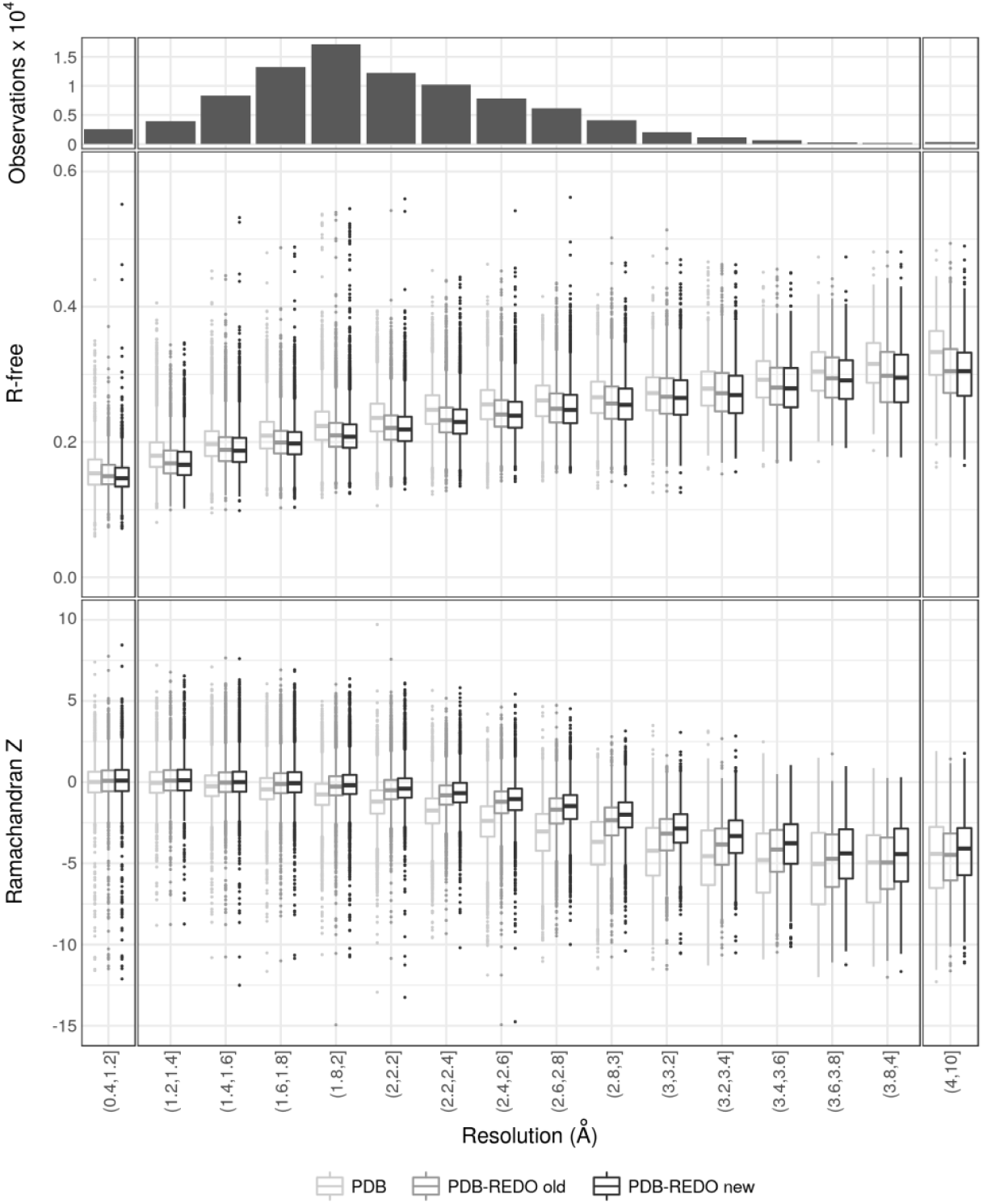
R_free_ and Ramachandran Z-score as a function of crystallographic resolution for entries present in PDB, in the PDB-REDO databank prior to the introduction of homology-derived H-bond restraints (PDB-REDO version 6.23), and in the PDB-REDO databank calculated with version 7.00. Outliers are shown when they are located beyond 1.5 times the inter-quartile range. R-free for PDB entries was determined by PDB-REDO for consistency.

Interestingly, the information source for homology-derived restraints can be analyzed in detail for every structure. In Figure 4 and interactive figures in Supplementary HTML, we show a directed graph to represent information transfer from any higher-resolution homolog to any lower-resolution homolog. About half of the PDB-REDO structures (nodes) in the network are connected (edges) to other structures, as they donate or receive H-bond restraints by satisfying the criteria for homolog use (see Online Methods). The connected graphs in the network typically correspond to protein families. The clusters have widely varying topologies and may be highly connected (Figure 4a) or may consist of a few structures that receive restraints from structures that only donate (Figure 4b). Interestingly, we found one single very large cluster, consisting of many smaller clusters connected to each other mainly by antibodies and lysozyme (Supplementary HTML). Using community structure detection^32^ we show that the modules in this graph also correspond to clusters of homogenous function (Figure 4c, 4d and Supplementary HTML). Visualizing and analyzing these clusters is an important tool for detecting likely changes within specific family members.

**Figure 4:**
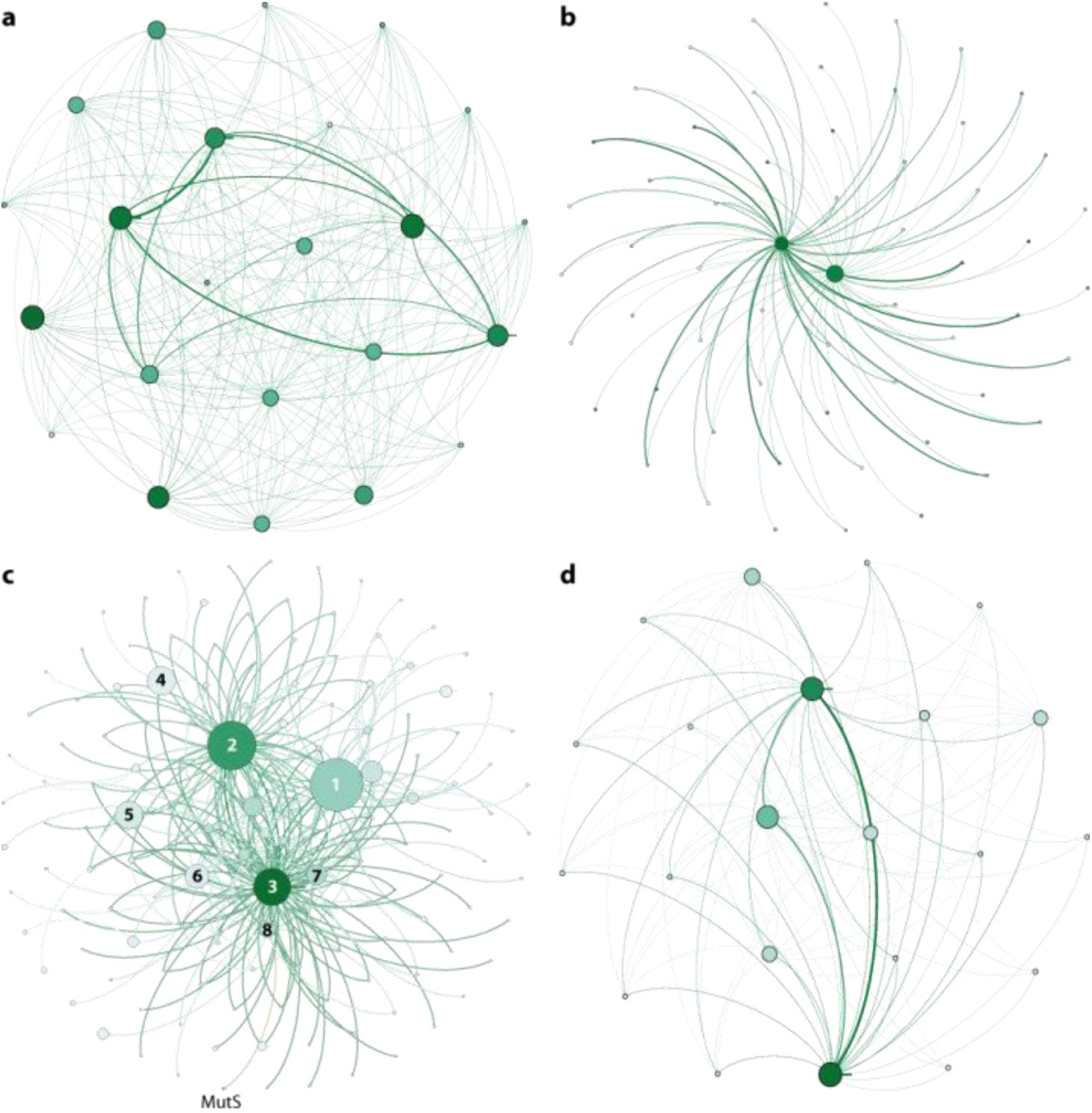
Network representations of H-bond information transfer between homologs. The nodes represent structures in the PDB-REDO databank. Node size and color correspond to the number of incoming edges and used resolution (darker is lower), respectively. The edge weight corresponds to the number of homologous chains. (**a**) Breast Cancer 1 (BRCA1). (**b**) Alcohol dehydrogenase (ADH). (**c**) Modules detected in the largest network. Node size reflects module size. The three most frequent terms in PDB TITLE records (stripped from English articles, punctuation, etc.) of the structure members in the labeled modules are 1) lysozyme, carbonic, anhydrase; 2) Fab, antibody, fragment; 3) antibody, Fab, HIV; 4) trypsin, inhibitor, thrombin; 5) HLA, peptide, class; 6) hsp90, bound, inhibitor; 7) ubiquitin, nucleosome, histone; 8) binding, maltose, bound. The MutS community (orange; MutS, mismatch, coli) is linked to community 8. (**d**) The MutS community.

The family of maltose transporters is an example where examining the cluster can aid analysis: the 3fh6^33^ structure in this family is receiving information from every other node/structure in this family (Figure 5b). Examining the structure in more detail indeed shows that the introduction of homology restraints has led to local improvements, *i.e.* better definition of the secondary structure content. The secondary structure for this protein became now more similar to the family (Figure 5a,c,d) in an unsupervised, automated manner, and would thus not mislead a potentially interested researcher to believe that this model is genuinely different to other homologues in secondary structure content.

**Figure 5:**
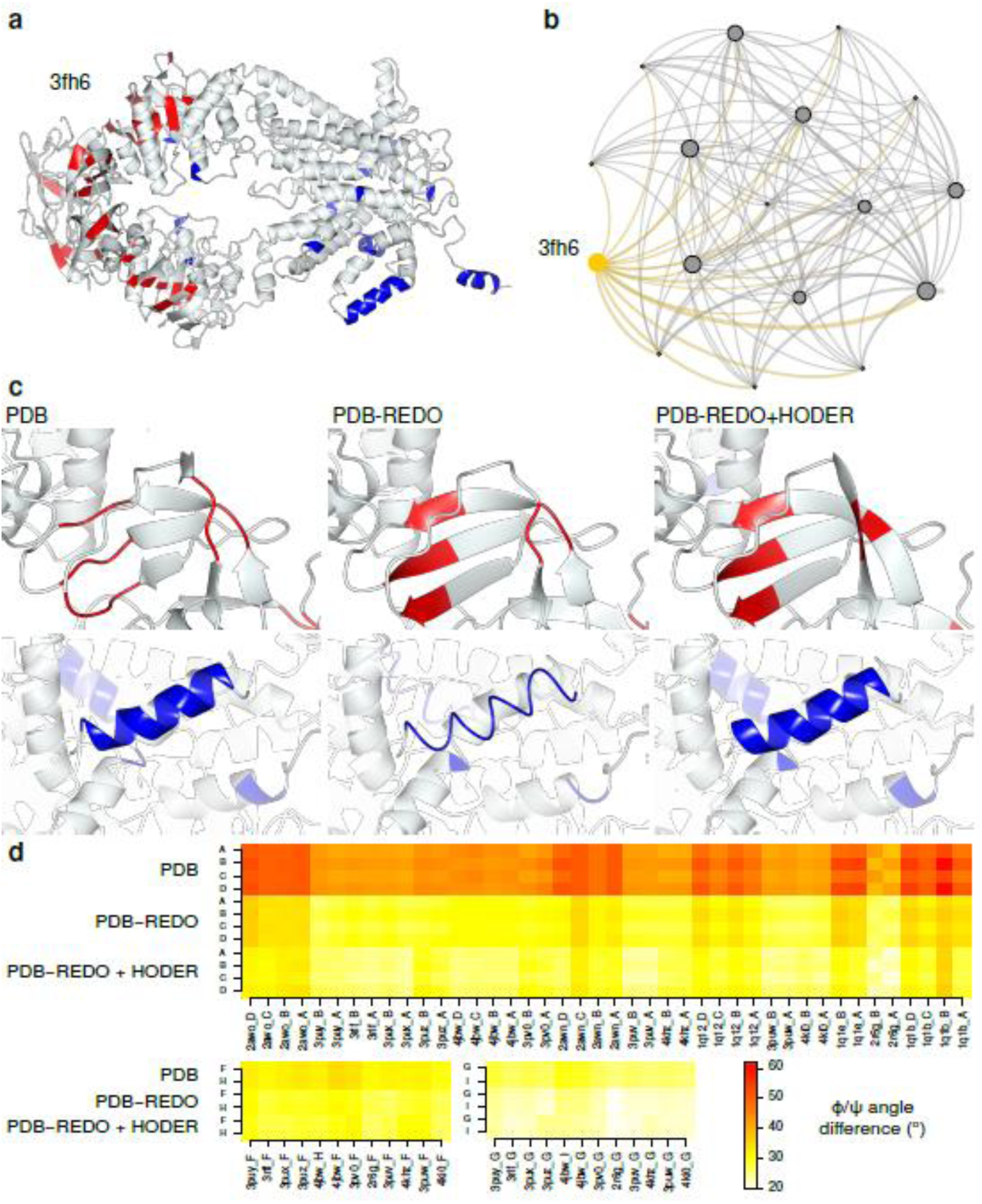
(**a**) The 4.5 Å structure model of E. coli maltose transporter (PDB entry 3fh6^33^) after PDB-REDO with HODER. All colored residues (strand in red, helix in blue) in the full structure are residues that changed in secondary structure between PDB, PDB-REDO and PDB-REDO with restraints from HODER. The secondary structure is best defined after using homology-based restraints. (**b**) The network neighborhood of homologous PDB entries that were used to define the restraints. The target entry, 3fh6, is shown in yellow. The node size corresponds to the number of incoming edges and edge thickness represents the number of homologous chains used. Small nodes are the high-resolution homologs that only donate information. (**c**, top) Details of a β-strand region are shown for PDB, PDB-REDO and PDB-REDO with HODER-generated restraints. The regularity of the strand is improved by PDB-REDO compared to the PDB and still further improved when restraints are used. (**c**, bottom) Details of an α-helical region in the same structure models. At such a low resolution, PDB-REDO requires the restraints from HODER to retain helical regularity. (**d**) The average absolute difference of φ/ψ torsion angles between 3fh6 chains and homologous chains for each homologous chain in the PDB, in PDB-REDO and in the new version of PDB-REDO with restraints from HODER. The chains A, B, C and D are homologous mixed α/β domains and there are two pairs of homologous α-helical domains: chains F and H and G and I, respectively. These three groups of homologous chains are shown separately. Especially the mixed α/β domains become much more similar to their homologous counterparts. All chains become still more similar to homologs when restraints are applied. Some homologs are clearly more similar in conformation to 3fh6 than others. All average angle differences fall in the range between 20 and 62 degrees presented in the legend.

Apart from making changes of local interest to specific structures, the new PDB-REDO databank can also provide a more reliable resource for data mining: for example, properties such as the Molprobity^34^ percentile and the ΔG of folding (calculated by FoldX^35^) are improved and become more uniform for protein families in PDB-REDO (Figure 6). Such uniform distributions can be much better learning sets for deriving empirical information by data mining protein structures, and can help improve modeling and analysis initiatives.

**Figure 6:**
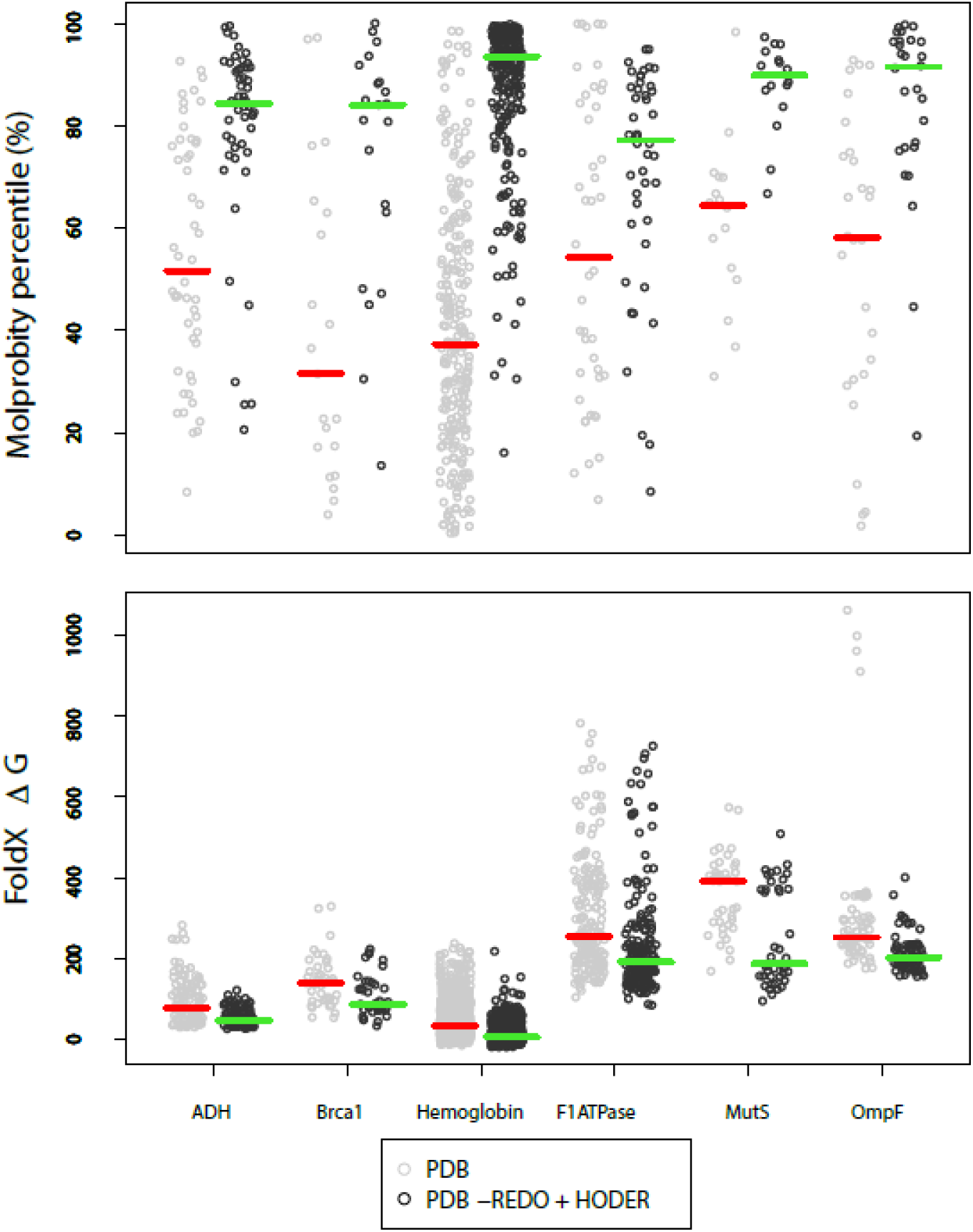
The free Molprobity^34^ percentiles (top) and energy (ΔG) of folding from FoldX^35^ (bottom) for each chain in the six investigated protein families. Data is shown for PDB and PDB-REDO with restraints from HODER. For the Molprobity percentiles, a single data point is shown per entry; for ΔG a score is shown per chain. The red and green horizontal bars indicate the median values.

## Discussion

Novel general and homology-based H-bond restraints targets, obtained by new algorithms mining the PDB-REDO databank, improve geometrical quality and the fit to X-ray data for low-resolution crystallographic structure models. This improvement often goes beyond the reach of current methods. In standalone refinements, homology-based restraints perform equally well to restraints based on general data-mining. Within the PDB-REDO pipeline, however, homology-based restraints perform better than general restraints.

A difficulty with many methods based on reference structures is that their performance is dependent on the similarity of the reference structure to the target structure model. For example, in the LORESTR pipeline^17^, different reference models are tested. In that approach, separate restraints are generated for each reference model and the refinement will adhere to those restraints that are closest to the current model. In such an implementation, more homologs lead to more restraints for a single distance, and therefore distances will be restrained to a target more similar to the current distance. This effectively prevents the target model from changing, likely explaining the authors’ observation that more homologs did not improve refinement. In our approach, all homologs are used to generate one or a few targets per interaction, and therefore more homologs only lead to a better definition of the restraint targets. Using our approach also mitigates user dilemmas on model choice: a comprehensive search of likely models or combinations is computationally very expensive, but using a subset of homologs makes model selection semi-arbitrary. The methods presented here have the advantage of using all homologous structure models, making them more computationally efficient and more robust to differences in homologs than methods based on a single reference structure model. Additionally, the width of the H-bond-length distribution is represented in the restraints, allowing regions with more structural variation to be less tightly restrained and vice versa. This information is not available if only one reference model is used.

We expect that the multi-homolog methods presented here will not work as well if random short-range atom pairs are restrained instead of H-bonds. Unlike random distances, H-bonds can be validated based on well-established geometric criteria. Therefore the selection of restraints is much more reliable, albeit smaller than with random distances. With this in mind, the restraints defined here for H-bonds could be extended to other intramolecular interactions in a protein, such as π-π-, cation-π-, and anion-π-interactions. Unlike H-bonds, more than two atoms are involved in such interactions, hence more than a simple distance restraint is necessary to improve their geometry. The framework for restraining plane stacking interactions (as a proxy for π-π-interactions) is available in Refmac5 and is used by the program LibG for nucleic acid restraints^36^. Detailed studies into the geometry^37^ and thermodynamics^38^ of these interactions can aid in inferring which geometric parameters are best restrained.

The application of H-bond restraints PDB-wide in a massively parallel manner using HPC resource has generated a new resource for the biology community: a PDB-REDO databank that incorporates homology information and is uniformly re-refined and re-built with a single software version. By eliminating software idiosyncrasies from the generation of the final structure model and by using homology information, structure similarity within a protein family can be analyzed optimally. That is, in PDB-REDO, differences between models are more likely to be true differences instead of refinement-related inconsistencies and the models are therefore more informative. Indeed, the PDB-REDO databank was previously used to systematically study OH cleavage from tyrosine^39^. Moreover, using a higher-quality structure data resource may prevent incorrect conclusions from dubious data. For example, a recent study^40^ detected a number of “novel zinc coordination geometries”, most, if not all, of which were simply errors in the input PDB data^41^.

Importantly, H-bond restraints are aimed at improving the geometry of protein structure models and are therefore not solely applicable to models solved by X-ray crystallography, but also to models obtained from cryo-EM. Models solved by cryo-EM still have a relatively low resolution compared to X-ray crystallography and often have homologous domains of higher resolution present in the PDB. H-bond restraints could also be applied to NMR and homology models but only once there is independent evidence that H-bonding partners are actually close; in these cases H-bond restraints should best be introduced at a final polishing stage of model optimization.

The new PDB-REDO databank is a valuable novel resource for two audiences. Structure-minded biologists can use the improved models to identify true features of particular structures in the context of a protein family. Bioinformaticians gain a resource that removes systematic errors from structural models to an extent that is likely to prove significant in the context of many types of global analyses, including homology modeling or automated feature analysis of protein structures.

## Availability

The methods discussed in this article have been incorporated into the PDB-REDO pipeline that is available as a webserver^3^ at https://pdb-redo.eu or as downloadable software. The new PDB-REDO databank is also available at https://pdb-redo.eu. The source code of HODER, *pdb2fasta*, and *detectHbonds* and the containerized versions of the entire PDB-REDO pipeline are available on request.

## Acknowledgements

This work is supported from the Netherlands Organization for Scientific Research (NWO) Vidi grant 723.013.003 to RPJ. Additional support was available from Horizon 2020 programmes West-Life (e-Infrastructure Virtual Research Environment project No. 675858) and iNEXT (project No. 653706). Computational resources and other support were granted by Johnson and Johnson.

## Author Contributions

BvB wrote all software modules discussed in this manuscript, analyzed data, prepared figures and wrote the manuscript; WGT containerized the PDB-REDO pipeline, deployed the PDB-wide PDB-REDO job, analyzed data, prepared figures and wrote specific methodology sections; MT participated in containerizing the PDB-REDO pipeline; GR and SS facilitated HPC implementation; GLG and JL contributed ideas, discussion and specific problematic PDB entries as metrics; RPJ wrote code integrating software modules in PDB-REDO; AP and RPJ designed the research program and finalized the manuscript; all authors reviewed the manuscript.

## Online methods

### Derivation, application and validation of general H-bond restraints

#### Derivation of hydrogen bonds

We have optimized our H-bond detection algorithm extensively, which lead to various cutoffs: unless specified otherwise, these values were empirically optimized to find the best set of H-bonds. We detect as H-bonds all pairs of potential donors and acceptors in a protein molecule that are distanced less than 3.5 Å away from one another. This results in a large initial set of H-bonds from which many will be filtered out after closer examination.

First, hydrogen atoms are modelled at the ideal positions. For donors that have more than one option for the ideal position of the hydrogen atom, such as the alcohols in Thr, Ser and Tyr, the best hydrogen is chosen as the one closest to the acceptor; others are deleted. McDonald and Thornton^19^ established criteria for the geometry of H-bonds; we work with slightly more lenient criteria (Figure 1). H-bonds whose geometry does not match the criteria are discarded.

Next, two general filters follow as a sanity check for each H-bond. First, H-bonds are filtered if the H-bond is physically blocked by another atom. Secondly, if the atoms in the H-bond are from different chains, the percentage of atoms with an occupancy not equal to 1.00 must be smaller than 80% for the H-bond to be kept (there are some instances in the PDB where quasi-multi-model refinement has been applied by simply generating two chains with the identical protein in a slightly different conformation; this filter prevents hydrogen bonding between those different “chains”).

After general filters, there are filters specific to main-chain and side-chain H-bonds. In the following, main-chain H-bonds are between two main-chain atoms; side-chain H-bonds are between two side-chain atoms or between a main-chain and a side-chain atom. The most important filter for main-chain H-bonds is a check for consistency with DSSP^20,21^: an H-bond is only kept if it is also a first-choice H-bond in DSSP. This immediately ensures that a main-chain donor does not form more than one main-chain H-bond, which is not allowed in our method because only one hydrogen atom can be donated. Additionally, main-chain H-bonds are only allowed if the donor and acceptor are at least three residues apart, and no H-bonds are allowed between donor and acceptor atoms that have different alternate codes.

Filtering of side-chain H-bonds is more elaborate because the diversity in H-bonds is greater. First, H-bonds between two alternates are removed when the D-A distance is smaller than 2.5 Å. Second, bifurcated H-bonds are deleted when they are superfluous. For instance, aspartic acid and arginine side-chains can form two H-bonds by bifurcation, but including cross-bifurcated H-bonds, up to four may be detected: the cross instances must then be deleted (Figure S3). Third, heavy atoms are only allowed to donate as many H-bonds as the number of hydrogen atoms they bind. Hence, alcoholic side-chains and backbone amides can only donate one H-bond while a lysine may donate three. Acidic side-chains usually lack a hydrogen and are therefore considered to be acceptors only. Fourth, no general target could be defined for some seven H-bond types (Figure S2) involving histidine because they are very rare: no restraints will be produced for those H-bonds. Finally, some H-bonds have both an ambiguous donor and acceptor and will be detected twice. The second instance is therefore removed.

When all other filters have been applied, conflicts can still arise between the set of main- and side-chain H-bonds. If a main- and side-chain H-bond are based on the same hydrogen atom, the bond with the longest H-A distance is discarded.

After applying all filters, there are still two types of side-chain H-bonds that we consider less likely than other H-bonds. Therefore, these H-bonds are only restrained with a homology-based target and not with a general one. The first case concerns H-bonds where donor and acceptor are in the same or sequential residues. This is uncommon and only allowed if the target can be derived from homologs. Serine, threonine, glutamic acid and glutamine-Oε form the exception to the rule, as they commonly bond to their own backbone^42^ and are therefore allowed to be restrained with general targets. Secondly, H-bonds are flagged from long side-chains of arginine, glutamic acid or glutamine if they only make a single side-chain H-bond. Such H-bonds are not as reliable because the conformation of long side-chains is often less ordered.

#### Fitting H-bonds distributions

The H-bond-length distributions were fitted with two-sided normal distributions using simplex optimization^43,44^ of least squares in combination with a linear weight on the height of the distribution. We give greater weight to the top of the distribution than to the edges because H-bonds in the middle of the distribution are more likely correct. However, the normal distribution was fitted on two sides of the mean separately because the respective variances were very different for most categories. The final target distance is then defined as the fitted top of the distribution and the standard deviation is set as the average of the standard deviations of the two sides. This was necessary because the Geman-McClure restraint penalty function in Refmac*5^28^* is symmetrical. We determined manually that distributions based on >100 observations can be fitted with reasonable accuracy. H-bonds of the rarest six types (out of 119) are currently not restrained. Since the target fitting routine is fully automated, the targets can be updated regularly and thus become more precise and complete when more data is available.

For choosing the weight of H-bonds in refinement, integral weight values of 1, 2, 3, 4, 6, 8 and 10 (ProSMART/Refmac5’s default value) were tested. The optimal restraint weight was determined to be 2.

#### Initial testing and validation of H-bond restraints

A program called *detectHbonds* was written, which takes a PDB file and generates H-bond restraints in Refmac5’s external restraint format^8^. The effect of the H-bond restraints was evaluated by running refinements twice per structure model, once with and once without restraints. Settings for geometric restraint weight, B-factor model, jelly-body restraint weight, twinning settings and B-factor restraint weight were taken from PDB-REDO runs with and without the use of our new restraints (the implementation in PDB-REDO is discussed in the section “Homology-based H-bond restraints in PDB-REDO”). All refinements were run for 50 cycles to ensure convergence.

The effect of H-bond restraints is greater at a lower resolution, because the effective restraint weight is larger and because there is generally more room for improvement. An initial survey showed that at a resolution better than 2.5 Å, the effect of H-bond restraints is negligible. Therefore, we only included structures of lower resolution in our test set. Five non-homologous entries (*i.e.* non-BLAST^23^ hits under default settings) were chosen at random with a resolution between 2.5 and 2.6 Å, then 5 entries between 2.6 and 2.7 Å, and so on up to the lowest resolution in the PDB. Seven very low resolution entries (5-8 Å) were removed from the test set because suitable refinement parameters for Refmac5^28,45^ could not be determined. An additional requirement for test entries was the presence of sufficient homologous chains in PDB-REDO, while the test entry itself had the lowest resolution among these homologs. This criterion was added because of the development of homology-based restraints which are discussed in the next section. The final test set contained 155 entries.

The difference between the models resulting from Refmac5 runs (version 5.8.0155) with and without H-bond restraints gives a direct measure for the influence of the H-bond restraints. We used R/R_free_ to measure crystallographic fit of the model to the data and use WHAT_CHECK^27^ (version 14) for geometrical validation. Specifically, we used WHAT_CHECK’s Z-scores for 1^st^ and 2^nd^ generation packing quality^46^, and rotamer torsion angle and Ramachandran plot normality (*i.e.* X_1_/X_2_ and φ/ψ normality). Also, WHAT_CHECK’s number of unsatisfied H-bond donors and acceptors and the number of atomic clashes or ‘bumps’ were taken into account in geometrical quality analyses. The statistical language R^47^ was used for calculation of statistics and plotting.

#### Improvement of crystallographic refinement using general H-bond restraints

H-bond restraints on the basis of general targets improve the refinement of low-resolution structure models in a large majority of cases (Figure S2, Table S3). Mainly packing and Ramachandran angles are improved, while marginal average effects are observed for R_work_ and R_free_. Upon adding H-bond restraints to refinement, the number of bumps decreased in 129 cases, remained equal in 12 cases and increased in 10 cases. The number of unsaturated H-bond donors and acceptors decreased in 84 cases, remained equal in 25 cases and increased in 42 cases. The only metric that deteriorates on average is side-chain rotameric angle quality.

The torsion angles of the backbone, φ and ψ, are decidedly improved, as is reflected in the Ramachandran Z-score (Figure S2, Table S3), in contrast to the side-chain torsion angles χ_1_ and χ_2_ (rotamer Z-score). Since the patterns of the main chain are generally more well-defined as those in side chains, it was expected that main-chain H-bond restraints would have more impact than side-chain restraints. Indeed, this is the case, as the restraint effect is almost as large when side-chain restraints are left out (Table S3). The well-defined patterns of main-chain restraints also explain why main-chain torsion angles are more easily improved: the side chains do not so strictly follow the patterns outlined by their restraints. Therefore, the restraints are less likely to help a side chain find its optimal orientation. Moreover, the standard deviation of side-chain restraints is generally larger and the target is thus less strictly adhered to in refinement.

#### Comparison restraints with ProSMART and *phenix.secondary_structure_restraints*

Our H-bond restraints lead to better results with Refmac5 than the restraints generated from ProSMART and *phenix.secondary_structure_restraints*^9^. There are two differences between all implementations: the H-bond selection and the target distances. The latter do not greatly influence performance, because replacement of the targets in our restraints by targets of ProSMART (2.8 Å) or Phenix (2.9 Å) only marginally changed the total performance in simple refinements. Therefore, the main reason for improved performance must be better H-bond selection. In Figure S4, the overlap between restraints for the different programs is shown. A large part of the restraints is shared between all programs. Phenix makes the fewest restraints, because only restraints for secondary structure elements are generated in their implementation (from definition by ksDSSP). Such restraints generally deliver the highest benefit per restraint and hence, the Phenix implementation works very well. However, that implementation misses out on the opportunity to gain extra performance by selecting a wider range of H-bonds. ProSMART does select a wider range of H-bonds, but unfortunately this selection also includes some distances that are better not restrained. In particular, we noticed a strong tendency of ProSMART to restrain both i,i+3 and i,i+4 main chain pairs in a-helices, where only the i,i+4 pairs should be restrained. This explains why especially the Ramachandran score is not improved by H-bond restraints from ProSMART: the application of conflicting restraints will distort the backbone rather than stabilize it. A better selection of H-bonds will therefore likely make H-bond restraints generated by ProSMART more useful.

#### Analysis and visualization of restraint satisfaction

A python script, distel.py, has been written to analyze the satisfaction of the restraints. The script matches each restraint to atoms in the corresponding PDB file and then calculates the distance between those atoms. The target and standard deviation of the restraint are used to compute a Z-score. Individual outliers of > 4.0 can be shown and the script computes a global rmsz score of restraint obedience. Additionally, a YASARA^48^ scene can be generated to visualize the restraints (Figure S5). The violation of restraints is also visualized by color-coding the arrows that indicate the restraints.

### Homology-based H-bond restraints

#### Derivation of protein sequence using *pdb2fasta*

The program *pdb2fasta* derives the sequence of a protein accurately from a PDB file. If available, *pdb2fasta* uses SEQRES records in PDB files to derive the sequence of unmodeled parts of the protein. Modeled and unmodeled residues are shown as uppercase and lowercase letters, respectively, as was done previously in the SEQATOMS tool^49^. In contrast to SEQATOMS, *pdb2fasta* does not use the annotation in the PDB’s mmCIF files because the annotation does not meet our requirements. For example, residues are labeled as modeled in mmCIF if just a single atom has been modeled, however, we require at least a complete backbone to look up homologous hydrogen bonds. Therefore, *pdb2fasta* also reads the ATOM records of PDB files in full and labels a residue as ordered only if all four backbone atoms are present. Unmodeled residues are found by mapping the sequence from the SEQRES records onto the ATOM sequence. As a special feature, *pdb2fasta* converts 73 types of prevalent modified amino acids to their base amino acid (e.g. trimethyllysine and selenomethionine are converted to lysine and methionine, respectively) to improve sequence alignments downstream.

The user can also specify an input FASTA file. The sequence from this file are treated as SEQRES records and thus mapped to the ATOM sequence. This allows the user to specify the unmodeled stretches manually. *Pdb2fasta* attempts to map each PDB chain to one of the entries from the FASTA file. An exact match with the ATOM records is immediately accepted; in other cases it aligns the sequences from the ATOM records and from the FASTA file. This can result in an alignment of all ATOM records to part of the FASTA sequence: in that case, the remainder of the FASTA sequence is assumed to be unmodeled and the alignment is accepted. It can also happen that there are several mismatches between the ATOM sequence and the FASTA sequence. If the sequence identity is over 90%, the match is accepted and the user is alerted to the mismatches. The sequence that is output is then the input FASTA sequence corrected for the mismatches found with the PDB file. In case both an input FASTA file and SEQRES are present, the sequence from the FASTA file is preferred (if it is accepted as a hit).

#### Selection of homologs

Homologs are then identified in a BLAST run against a sequence database containing only PDB-REDO entries. By default, BLAST hits are selected as a homolog if the sequence identity is at least 70% and the E-value is smaller than 10^-3^. Homologs with a resolution worse than the query are discarded. Fully optimized PDB-REDO coordinate files are retrieved for each homolog. HODER reads these PDB files and maps the residues of the BLAST hit to those of the query as in the BLAST alignment. Users can also add their own PDB files of homologs to HODER, which will be used as extra homologs provided they pass the sequence similarity criteria. If, for instance, one is working on a series of ligand soaks, this feature enables them to make full use of their homologous data in refinement before any models are made publicly available.

#### Deriving restraint target values from homologues

After mapping homologous residues onto query residues, the distances of an interaction in all homologs are computed. To calculate this distance, the equivalent homologous atoms must be found. If the amino acid types are the same in query and hit, the homologous atoms are usually the same atom types in the homologous residues. An exception can occur if the residue contains equivalent atoms. For example, if the acceptor of a certain H-bond is an aspartic acid’s Oδ1, the homologous atom could be the Oδ1 or Oδ2 in the homologous aspartic acid. The homologous atom is then selected by picking the atom whose torsion angle most closely resembles that of the atom in the query.

If no homologous atom can be identified for certain H-bond, that homolog will be skipped for that H-bond. However, even when the amino acid types do not match between query and homolog, homologous atoms can often still be found. For main-chain H-bonds, the homologous atom is simply the same backbone atom in the homologous residue. The backbone nitrogen of the imino acid proline is the only exception, because it has no hydrogen atom bound and hence cannot donate an H-bond. For side-chain H-bonds, homologous atoms can only be identified if the homologous residue contains an equivalent atom, which is the case for Asp and Asn, Glu and Gln and Ser and Thr. For instance, an asparagine contains an Oδ1 that can be the homologous atom for an aspartic acid’s Oδ1 or Oδ2. Such atoms are only assigned as homologous atoms if the torsion angle differs by less than 90° from the query atom.

A final complication in assigning homologous atoms can arise if the homologous residues contain alternate conformations. The rotamers of the alternates of the homologous residue are then evaluated and only if exactly one alternate, with an occupancy greater than 0.25, matches the rotamer of the interaction, the homologous atoms are assigned from that alternate.

Aside from actually being able to find homologous atoms for both donor and acceptor of an H-bond, we apply two other filters to ensure we find homologous distances of high quality. First, the fit to the electron density for both hydrogen bonding partners, represented as the per-residue real-space correlation coefficient (RSCC)^50^ obtained from EDSTATS^51^, should be sufficient. By default, the worst 3% of residues in each structure model are ignored. Second, for side-chain interactions it is demanded that the homologous atoms are in the same rotamer as the query atoms. Rotamers are considered different if one of the X angles along an sp3-sp3 bond changes by at least 50°, or if a χ angle along an sp^2^-sp3 bond (i.e. in Asp, Asn, Glu, Gln, Phe, Tyr, and His), changes by more than 30°. If needed, Asp, Glu and Tyr side-chains are flipped to minimize the change in torsion angles.

The distances computed from all homologous H-bonds are subjected to 1D *k*-means clustering^25^ after filtering out distances longer than 6.0 Å. A maximum of three clusters is fitted to each interaction; the optimal number of clusters is determined by the Bayesian information criterion^26^. When a limited amount of homologous distances is available (< 13 data points), the maximum number of clusters is reduced to two. Clusters are discarded if they contain less than 5 distances or less than 25% of the total number of observations. Subsequently, distances are removed from clusters if they are an outlier according to Grubbs’ outlier test^52^ with a 90% confidence limit and deviate at least 0.25 Å from the cluster mean. If targets for the same interaction differ by less than 0.1 Å, the clusters are merged and a new target is defined. Finally, if all targets for a given interaction are longer than 3.5 Å, the maximum allowed length of a H-bond, no restraints are written because the H-bond is likely incorrect. All values mentioned here were chosen based on observations of corner cases in the data.

### Testing homology derived restraints

We obtained a considerable improvement of the geometry of protein structure models (Figure S6, Table S4) in our test set, but overall improvement achieved by using homology-based restraints is similar to that of general restraints (Table S4). Packing is improved somewhat less; torsion angles are improved somewhat more. Importantly though, we find far fewer cases where the backbone angles are deteriorated. The torsion angle improvement can be explained by the fact that weak H-bonds in the main chain will no longer be restrained to a short distance at which they slightly deform the backbone. Similarly, the packing score will likely improve less because tight backbone is not enforced as much as in restraints based on general data.

#### Homology-based H-bond restraints in PDB-REDO

The new tools to generate additional restraints were added to the PDB-REDO pipeline. *Pdb2fasta* is run in the early stages of the process after the program *extractor*^1^ if that program establishes that the input model contains protein. If there is a user-provided FASTA file, any conflicts between that and the atom records of the input PDB file are reported to the user. If the resolution of the diffraction data is 2.8 Å or worse, or if there are fewer than 2.5 reflections per heavy atom, homology restraints are created automatically after the initial calculation of crystallographic R-factors. This behavior can be repressed, or at higher resolution enforced, from the command line. With the extracted FASTA sequence, *BlastP*^23^ is run against a sequence database that contains only entries of the PDB-REDO databank. Next, the output of the BLAST search is fed to *HODER* which automatically downloads suitable homologous structure models from the PDB-REDO databank or uses the corresponding files in a local copy of the databank. User-provided homologous structure models are also included at this stage. Users can specify at the command line if *detectHbonds* should be run instead of *HODER*. The restraint*s* written by *HODER* or *detectHbonds* are added to all subsequent Refmac5 calculations. After the first model re-refinement and model rebuilding, *HODER* or *detectHbonds* is run again to update the restraints for the new model coordinates.

*distel.py* is run after the re-refinement (against the restraints at that stage of the procedure) and at the final stages of PDB-REDO with the final restraints.

#### Assessing structural differences between homologues

Homology-based restraints work decidedly better than general restraints in the PDB-REDO pipeline. The packing is improved for homology-based restraints by almost as much as general restraints while the increased performance of homology-based restraints for the torsion angles is more pronounced than for single refinements. In addition, we obtain an additional improvement in R_work_ and R_free_ that was not observed for single refinements. The R_free_, which was previously improved by PDB-REDO by 2.2% on average, is now improved by 2.5%. Expressed in Z-scores, this additional improvement is 0.30 ± 0.11 (average ± standard error of the mean), which is similar to the improvements attained for most geometrical criteria. A difference between the single refinements and PDB-REDO runs is that additional restraints such as local NCS restraints^14,28^ and metal geometry restraints^30^ are used in PDB-REDO: the combination with these restraints apparently gives a boost to the performance of H-bond restraints. Also, PDB-REDO uses a two-step approach where the restraints are redefined after the first refinement phase. This means that the selection of restraints can be improved after an initial change of the structure model.

For defining the families we used the homology criteria as implemented in HODER to group structures that have at least one homologous chain to a family. A noteworthy observation upon inspection of structural differences between PDB-REDO protein families the φ/ψ torsion angles between homologous residues in the protein structure families to obtain an overall average root-mean-square fluctuation (rmsf), was that the rmsf was already decidedly lower in PDB-REDO than in the PDB before homology-based restraints were applied. Because homolog structures are often solved by molecular replacement, one might expect that they are biased in similarity towards one another. However, these results show this is generally not the case, as applying the PDB-REDO pipeline without any influence from homologous structure models actually makes protein structure family members more similar. Refining structure models with a consistent protocol therefore already removes some of the model differences that are not clearly supported by the experimental data.

#### Massively parallel computing for a novel PDB-REDO databank with homology information

The PDB-REDO core software components and third-party dependencies are under active development and new versions introducing new features are released frequently. PDB-REDO entries are generated for new Protein Data Bank entries on a weekly basis. The renewal of old PDB-REDO entries, however, is limited by our in-house computational resources. As a consequence, at any time only a small fraction of the databank reflects new functionality (Figure S7). The availability of homology-based hydrogen-bond restraints was a strong motivation to regenerate the entire databank with PDB-REDO version 7.00, which we estimated to require about half a million CPU hours. We therefore decided to employ High Performance Computing (HPC) systems.

#### Containerization

The PDB-REDO computing pipeline consists of over 50 independent software components. Traditional installation of the PDB-REDO pipeline within the existing HPC system environment was hampered by system library restrictions and system package (dependency) issues. Recently the use of containers has become popular in software development. Emerging technologies like Docker (www.docker.com) and Singularity^29^ allow deployment of a consistent environment within a container that leverages host resources with minimal overhead (compared to Virtual Machines, for example). Furthermore, these technologies are highly scalable and suitable for HPC environments and scientific settings where reproducibility is essential^29, 53, 54^. We decided to containerize PDB-REDO to overcome the installation issues without sacrificing computation speed.

The installation of the most recent PDB-REDO and all its dependencies into a Docker images was automated. Every image is tagged with a unique identifier and labeled with versions of all software components, allowing versioning and archiving PDB-REDO results. The Docker images were converted to Singularity images using docker2singularity (https://github.com/singularityware/docker2singularity). Docker and Singularity images with and without licensed components YASARA^48^ (www.yasara.org) and FoldX^35^ are available from the authors upon request.

Automated image creation not only facilitated rapid in-house testing but also simple, fast and robust deployment of the PDB-REDO pipeline on any host supporting Docker or Singularity. The cross-platform portability of images guarantees reproducible results and eliminates maintenance tasks since it is easier to ship a new image than to update all PDB-REDO dependencies. Containers instantiated from an image can be executed like a normal PDB-REDO executable on any host that supports Docker or Singularity.

#### Deployment

The availability of the PDB-REDO image greatly simplified deployment on the San Diego Supercomputer Centre (SDSC) HPC systems Gordon and Comet. PDB-REDO version 7.00 was run with Singularity version 2.1 for 101,119 jobs in single-threaded mode to maximize the embarrassingly parallel throughput on the 192 compute nodes (3072 cores) on Gordon. The 451 entries with more than 350,000 reflections were run in parallel mode on Comet’s large memory nodes where 460 GB of memory was available to every job. Fully optimized PDB-REDO structure models available d.d. 2017-02-03 were used as homology-derived H-bond restraints sources. These data were staged on local SSDs of every compute node for speed and robustness. The output from each job was written to the parallel Lustre filesystems on Comet and Gordon.

#### Run times

The elapsed wall clock time for the Gordon jobs is shown in Figure S8.

